# The ecology of cancer differentiation therapy

**DOI:** 10.1101/853002

**Authors:** Ricard Solé, Guim Aguadé-Gorgorió

**Affiliations:** ICREA-Complex Systems Lab, Universitat Pompeu Fabra, 08003 Barcelona, Catalonia, Spain; Institut de Biologia Evolutiva (CSIC-UPF), Psg Maritim Barceloneta, 37, 08003 Barcelona, Catalonia, Spain; Santa Fe Institute, 1399 Hyde Park Road, Santa Fe NM 87501, USA

## Abstract

A promising, yet still under development approach to cancer treatment is based on the idea of differentiation therapy (DTH). Most tumours are characterized by poorly differentiated cell populations exhibiting a marked loss of traits associated to communication and tissue homeostasis. DTH has been suggested as an alternative (or complement) to cytotoxic-based approaches, and has proven successful in some specific types of cancer such as acute promyelocytic leukemia (APL). While novel drugs favouring the activation of differentiation therapies are being tested, several open problems emerge in relation to its effectiveness on solid tumors. Here we present a mathematical framework to DTH based on a well-known ecological model used to describe habitat loss. The models presented here account for some of the observed clinical and in vitro outcomes of DTH, providing relevant insight into potential therapy design. Furthermore, the same ecological approach is tested in a hierarchical model that accounts for cancer stem cells, highlighting the role of niche specificity in CSC therapy resistance. We show that the lessons learnt from metapopulation ecology can help guide future developments and potential difficulties of DTH.

## I. INTRODUCTION

Cancer is a set of complex diseases, and the success of tumor progression (and the eventual death of its recipient organism) requires a number of changes to make cells capable of overcoming selection barriers. These changes provide the source of proliferative power that makes tumors able to expand and evolve (Weinberg 2015). One particularly remarkable feature of cancer cells is the loss of molecular markers associated to the differentiated state. As the tumor evolves, some cancer cells appear to be in a de-differentiated state closer to early developmental stages, similar to that of normal stem cells, with increased potential for self-renewal and plasticity (Magee et al 2012). To some extent, cancer is a disease of multicellularity: the cooperative order required to maintain organism’s coherence is broken in favor of unicellular-like traits (Aktipis et al 2015, Davies et al 2011).

The standard treatment of tumors has been grounded in the use of either specific cytotoxic drugs or radiotherapy, or a combination of both. The success of this approach has been discussed and even questioned over the last decades (Gatenby 2009). Treatments involving a general mechanism of cell damage associated to toxicity are often inefficient and can trigger evolutionary pressures that select aggressive and resistant clones (Pepper et al. 2011). As a consequence, cytotoxic therapies can create undesirable side effects such as the development of metastasis. To a large extent, despite the undeniable success in our increasing understanding of the underlying molecular basis, cancer remains incurable. Because of these limitations, novel approximations have been proposed mainly from evolutionary and mathematical biology. They are based on the view of cancer as an ecological and evolutionary problem (Merlo et al 2006; Korolev et al 2014). In particular, ecological principles can guide alternative insights to cancer development and treatment (Basanta and Anderson 2013).

One specially promising alternative to conventional cytotoxic agents is the use of so called *differentiation therapy*. Here the approach, early suggested more than 50 years ago (Pierce and Wallace, 1971; Pierce 1983) is inspired in the observation that one hallmark of cancer is the loss or blocking of differentiation that leads to cells with increased potential for self-renewal and plasticity. Differentiation therapy (DTH) involves the use of diverse molecular agents able to induce differentiation in cancer cells. Since differentiated cell types are a terminal branch of development, the goal is to facilitate this process and remove cancer cells from the proliferative compartment. A growing family of DTH agents include neural growth factors, all trans retinoic acid, arsenic trioxide, butyric acid or cAMP, which have been shown some degree of differentiation-inducing capability both in vitro and/or in vivo (de Thé 2018). The success of DTH is well illustrated by the best known case study, namely its use in Acute Promyelocytic Leukemia (APL) by means of a combined cytotoxic therapy with all-trans retinoic acid (RA) (Huang et al., 1988).

A few numbers reveal some features of the impact of DTH. Again within the context of APL, before the use of DTH, cytotoxic-based therapies increased the likelihood of remission from 50 to 80 % but with only a third of long-term survival. The combination with RA changed drastically the situation, with 90 % remission and a 75 % cure (see Dela Cruz and Matushansky 2012 and references therein). Interestingly, when DTH alone is used, despite cell differentiation perfectly well identified (it can actually be massive) only combination with standard cytotoxic agents seems to account for long-term disease remission (de Thé 2018).

Over the last years, DTH agents have been also used for treating solid tumors. In contrast with the APL case study, the therapeutic effect of the differentiation-inducing agents on solid tumors is not strong when compared with that of conventional chemotherapeutic agents. However, because most of the differentiation-inducing agents can potentiate the effect of conventional chemotherapy or radiation therapy, combination therapy might be used as a second- or third-line therapy in patients with advanced cancer. Are the solid nature of the tumors, their genetic complexity or their hierarchical architecture leading factors for this limited success? Here too, a theoretical model can be helpful in interpreting the role of spatial competition effects and niche specificity in understanding the possibilities of DTH. The analysis of how differentiation therapy modulates the cancer habitat is based in an ecological approach to tumor dynamics inspired in well-established results from habitat loss and fragmentation in metapopulations (Mollanen and Hanski 1998, Hanski 1999).

## II. METAPOPULATION MODEL OF TUMOR DIFFERENTIATION THERAPY

The simplest mathematical approach taken here is based on the assumption that two different therapies act together on the growth of a cancer cell population. As a first dynamical description able to characterize a wide set of disease types we use a generic logistic growth model (Gatenby and Gillies, 2008), where cellular replication saturates as the tumor population approaches the carrying capacity *K* of the micro-environment. Sigmoidal growth has been proven to capture the tumor microenvironment effects of spatial constraints, resource limitations or intercellular growth inhibition in a wide range of malignancies such as breast cancer (Norton 2008), colorectal cancer (Misale et al. 2015) or chronic lymphocytic leukemia (Gruber 2019) The cancer cell population is also inhibited in two different ways. The first corresponds to standard therapies, based on cancer-targeted cytotoxic drugs. In this first scenario, a population of cancer cells *C* follows:

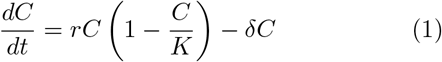

where for simplicity the carrying capacity will be normalized to one (*K* = 1) and thus *C* can be understood in terms of the fraction of host tissue occupied by the tumor. The last term in the rhs indicates the linear decay caused following from cellular death rate *δ* that can be increased under chemotherapy. This is equivalent to the well-known Levins model, where growth and decay would be related to colonization and extinction (Levins 1969). The analysis of this system reveals that two equilibrium states *C*\* are possible: extinction *C*\* = 0 and 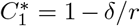. Tumor growth will occur when *r* > *δ*, i. e. if growth overcomes the negative impact of treatment.

How can differentiation treatment be introduced in this approach? The impact of DTH is dynamically very different. Previous research has studied mathematical modeling of tissue hierarchies by introducing differentiation as a rate at which progenitor cells transition into noncycling phenotypes (Dingli and Michor 2006, Werner et al. 2016). However, before getting into this multi-layer approach (Fig. 1b), we aim at understanding a specific effect of DTH that has not been included in previous models: what is the effect of non-cycling, differentiated cells as they occupy the habitat that the tumor needs for expansion? (Fig. 1b, dashed box). Lessons from habitat fragmentation have shown that these effects might be potentially key in extinction of colonizing species (Mollanen and Hanski 1998, Hanski 1999). In this context, the DTH scenario discussed below is inspired in the nonlinear behavior how fragmented landscapes (fig. 1c). Reductions of habitat size, along with stochastic death, define viability thresholds that we will connect with tumor extinction dynamics. In particular, these models reveal a somewhat counterintuitive feature that is relevant to our paper: a reduction of habitat does not simple imply a similar population reduction. Inste ad, extinction can occur despite the existence of a remaining habitat.

**FIG. 1:**
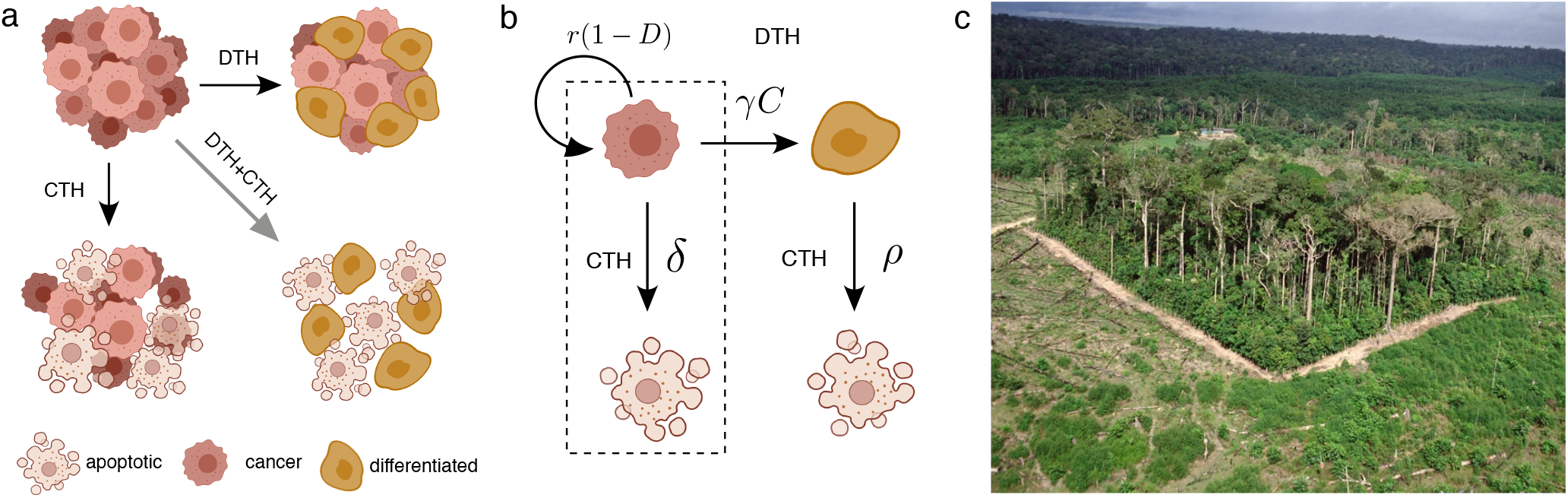
A metapopulation model of tumor differentiation therapy. In tackling alternative treatments to cancer progression, DTH exploits the potential of blocking tumor growth by activating differentiation pathways. A combination of cytotoxic therapy (CTH) and differentiation therapy (DTH) as shown in **(a)** can successfully kill the tumor when Both separately cannot. In **(b)**, both the complete model including the dynamics of the differentiated compartment and the minimal habitat-loss approach (dashed box) is shown. The DTH+CTH model is inspired in studies of habitat fragmentation **(c)**, where habitat reduction, along with stochastic mortality, can trigger species extinction. Drawings were made with Biorender.

As a fraction of cancer cells gets differentiated, they have an impact in population dynamics as they contribute to the overall population and thus limit the potential carrying capacity of the system. Studies on control networks indicate that not only spatial or resource constraints but also cellular signaling contributes to these dynamics (Vainstein et al. 2012, Yang et al. 2017, Komarova et al. 2018). If *D* weights the effectiveness of the DTH the simplest extension of the previous model incorporates the amount of

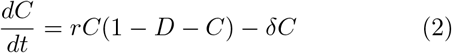

The ecological equivalent here is the extended Levins model incorporating habitat loss (Bascompte and Solé, 1996, Figure 1b, red box). In habitat loss models, the *D* term is associated to the amount of habitat that has been degraded thus being unavailable to colonization. The interpretation within the context of DTH is easy: the fraction of cancer cells that have become differentiated introduce a shift 1→1 − *D* in the maximum available habitat.. The non-trivial fixed point is now:

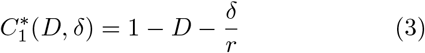

In this case, cancer decay will be expected provided that the fraction of differentiated habitat (and thus the efficiency of DTH) is larger than a critical value:

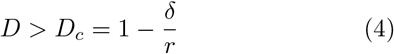

One particularly relevant and non-obvious result is that even if there’s apparently room for further growth, the dynamics of the system reveal a transition from cancer growth to cancer decay. Once the critical point *D_c_* is reached, tumor dynamics faces extinction.

In figure 2a we show a diagram for *D* against *δ* where the critical line *D* = *D_c_* has been used to separate the two phases associated to cancer progression and cancer decay. The lower axis indicates the efficiency of single cytotoxic therapy in the absence of DTH. A threshold is found for *δ_C_* = *r* as defined from model (1). By adding the second axis (differentiation) we can see that lower levels of chemotherapy are required to achieve tumor decay. This is at the core of our explanation for the success of DTH: the combination of both treatments can successfully achieve remission when the right combination of chemotherapy and differentiation is used. Since toxicity can be reduced provided that *D* is large enough, the diagram supports the observed success and long-term remission in APL. On the other hand, the levels of differentiation that are required for small *δ* can be very large (perhaps unrealistically large). An important point needs to be made here: could DTH only also achieve remission? The model in this case reads:

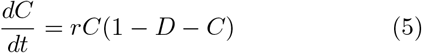

which can be shown to behave always in the same way: a logistic growth towards an intermediate level *C*\* = 1 − *D* with no threshold value. This implies, and seems consistent with clinical evidence, that DTH alone will fail to succeed given the lack of a remission threshold.

**FIG. 2:**
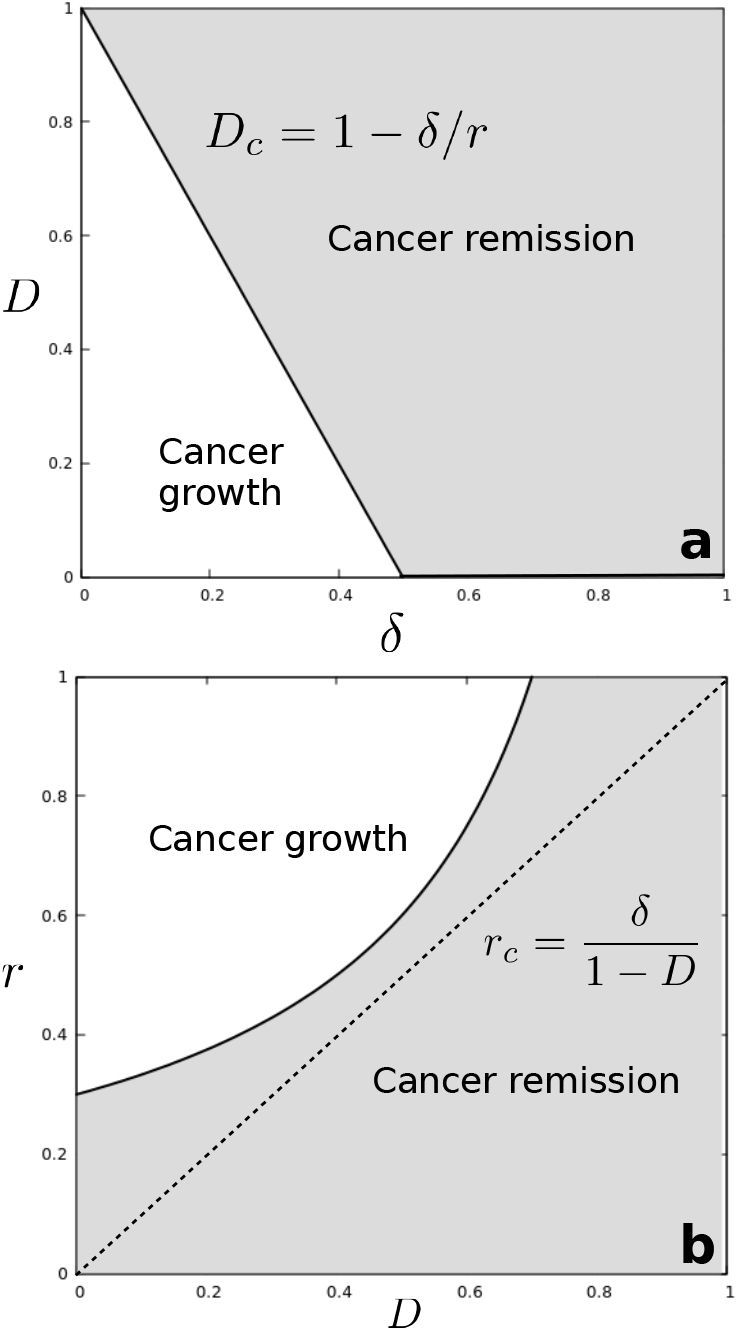
Phase space of cytotoxic-differentiation combination therapies. In **(a)**, the use of DTH can induce cancer remission even for death rates smaller than the tumor cells replication rate. In **(b)**, the nonlinear effect of DTH is pictured. For *C* ⪡ 1 − *D*, the replication rate necessary for cancer outgrowth *r_c_* grows linearly with *δ* (dashed line), while increasing *D* imposes a stronger condition (curved black line).

We have used a specific functional form for population growth, and the previous analysis deals with steady states involving large populations. However, our results are robust, as shown by looking at early tumor growth in a differentiated environment, where *C* ⪡ 1 − *D*, provides interesting differences between the dynamical impacts of cytotoxic therapy and the possibilities of modifying environmental carrying capacity through DTH. The model described by equation (2) now reads

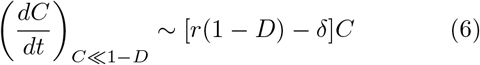

with exponential growth solution, i. e. starting from an initial population *C*(0),

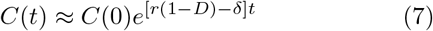

which gives cancer expansion only if the growth rate of the cancer cells is larger than a threshold value *r_c_*, namely

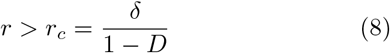

We can appreciate here the difference between the impact of cytotoxic therapy (acting linearly) and DTH (acting in a nonlinear fashion). In figure 2b we summarize these results by displaying cancer growth rates against the efficiency of DTH. In the absence of DTH, the tumor will grow if *r* > *δ*, but increasing habitat differentiation results in a nonlinear increase of the proliferation rate that tumor cells need to survive. A quick comparison with standard epidemiology models shows that this corresponds to epidemics suppression through vaccination: as more individuals are vaccinated and thus moved out from the pool of potentially infected individuals, the pathogen requires an increase in infectivity that might not be achievable.

How robust are the previous results? The model described above lacks an obvious dynamical layer: the amount of differentiated habitat follows in fact from cancer cells that have become differentiated. This can be clearly described by introducing an explicit, dynamical compartment of differentiated cells:

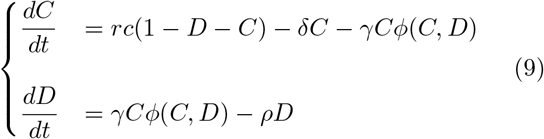

where *γ* is now the rate at which cancer cells differentiate, and *ϕ*(*C, D*) describes the possible control network relating differentiation to the amount of populations at play (Vainstein et al. 2012, Yang et al. 2017, Komarova et al. 2018). Furthermore, differentiated cells appear as they are produced by *C*, and disappear at rate *ρ* due to cellular death and removal (efferocytosis).

Cellular fate decisions, and differentiation in particular, are known to be controlled by regulatory signals related to different cell populations (Yang et al. 2017). However, the exact control networks governing differentiation processes are widely unknown. Knowing the clinical particularities of differentiation therapy, namely its major failure when administered as a single anticancer drug, can we infer possible expressions for *ϕ*(*C, D*)?

Among the possible functional forms for differentiation control *ϕ*(*C, D*), we expect a minimal function that is, at least, *f* (*C*). If not, stability of the (0, 0) attractor would follow from

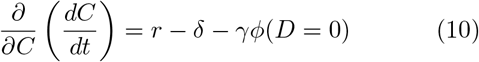

In this case, DTH alone would be able to eradicate tumor growth without chemotherapy (*δ* = 0), provided that

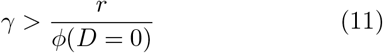

which contradicts, as discussed above, clinical observations (de Thé 2018). Our minimal assumption, consistent with computational results in (Vainstein et al 2012) is that regulation of differentiation is orchestrated by the amount of surrounding *C* cells, not considering secondary regulation effects by *D* cells:

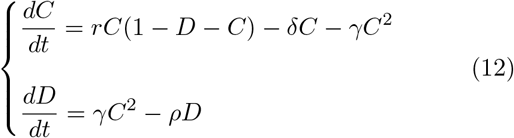

namely, differentiation increases due to cell-cell interactions, as introduced by the term *γC*^2^. This self-regulation term, similar to the one captured by a cellular automata in (Vainstein et al. 2012), is needed to capture that a larger differentiation rate alone will not be able to stop tumor progression (Fig. 3a). This assumption relies on a phenomenological approach to modeling, as the precise regulatory mechanisms of cellular fate remain largely unknown (Yang et al. 2017).

**FIG. 3:**
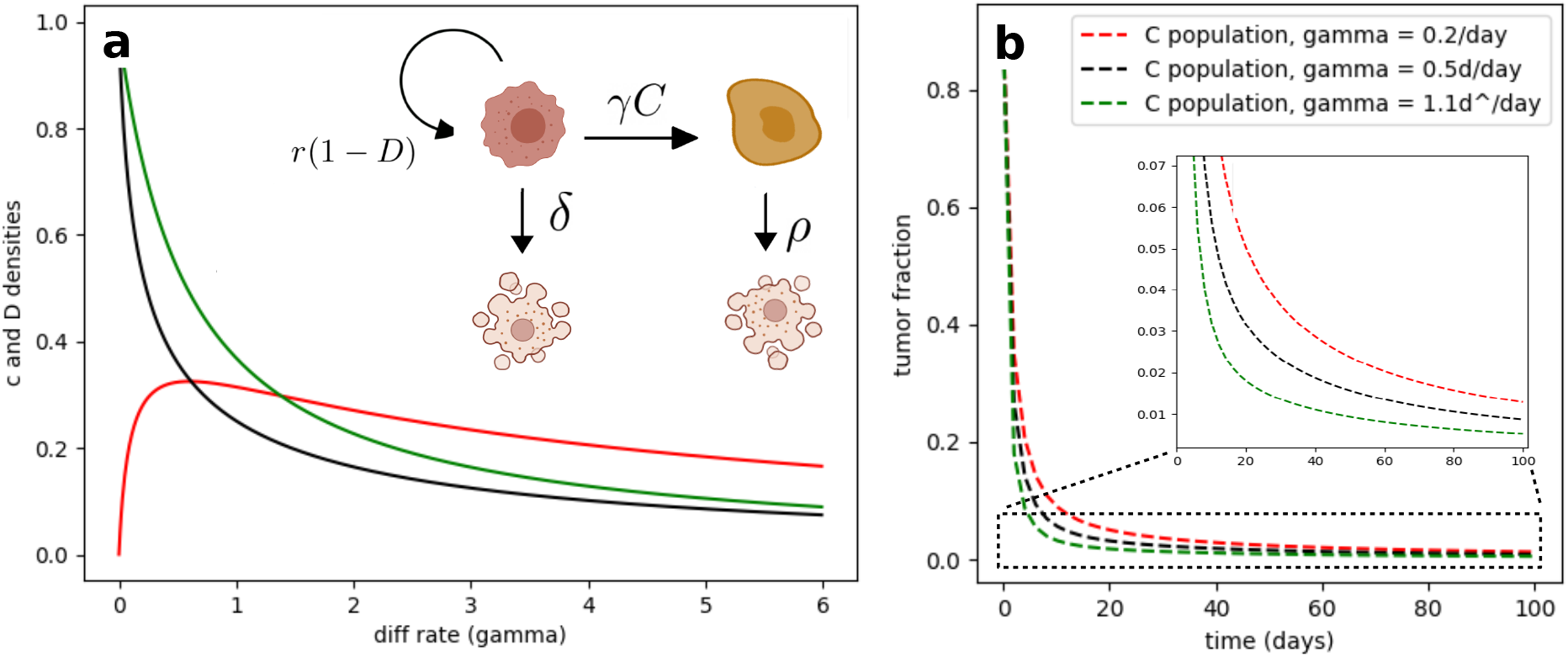
DTH and chemotherapy in the *C, D* model. In **a**, increasing levels of *γ* do not totally eradicate the tumor, but reduce the overall *C* (dark line) *D* (red line) populations, as it can be seen by computing the analytical attractor states (see SM). The green line indicates how the equilibrium *C* population would evolve in the absence of habitat-loss effects, indicating the relevance of considering the ecological dynamics of DTH. This is indicative of the possibilities of DTH when administered with cytotoxic therapies. In **b**, three different values of increasing *γ* show that a tumor targeted by CHT needs shorter chemotherapy interventions as DTH increases.

In this model, stability of the cancer-free state is only controlled by chemotherapy, provided that *δ* > *r*. What is the role of DTH here? This can be seen by obtaining the attractor states as a function of *γ*:

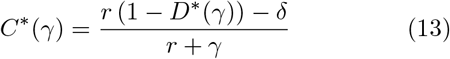

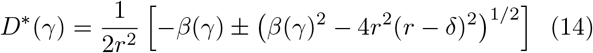

with

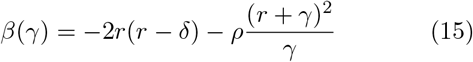

Here, modifying *γ* alone cannot modify the existence of at least one attractor state that is real and positive (see Figure 3a). However, because of the combined effect of differentiation in both *γ* and tumor growth constraints *r*(1 − *D*), agents incrementing *γ* will reduce overall tumor size, thus reducing the time and dose of a secondary cytotoxic agent needed to completely erase the tumor (Fig. 3b).

Several interesting points arise from describing cancer differentiation under the perspective of habitat-loss ecological models. First of all, the minimal model of DTH as habitat loss captures the existence of a differentiation threshold for tumor arrest. This highlights the role of habitat in tumor growth, with differentiation therapy driving an all-or-none response similar to that seen in APL treatment outcomes (Huang et al. 1988). When introducing the cellular dynamics of the differentiated compartment *D*, the sharp threshold gets diluted, but the model still shows the opportunities of DTH provided it is not delivered as a single agent.

Both model versions indicate that DTH is only effective when combined with cytotoxic therapies directly targeting the cellular death rate *δ* (Figure 1). This could explain why arsenic, that triggers p53-driven senescence apart from differentiation (Ablain et al. 2014), is functional as a single-agent therapy, while retinoic acid and other differentiation drugs that do not target cell death specifically require combined cytotoxic therapy to success (Dos Santos et al. 2013). The study of differentiation and tumor habitats becomes even more relevant in the context of Cancer Stem Cells (Meacham and Morrison, 2013). How do results change when a tumor seeding population resides in a different habitat?

## III. DIFFERENTIATION THERAPY IN HIERARCHICAL TISSUES

A broad range of cancerstypes are hierarchically organized, with a population of *cancer stem cells* (CSC) driving tumor growth and plasticity (Meacham and Morrison, 2013). Besides the relevance of this in radioand chemotherapy resistance (see e.g. Dean, Fojo and Bates, 2005), we are interested in understanding if the hierarchical architecture specific to a stem cell compartment is related with the fact that most solid and genetically complex tumors do not show valuable responses to differentiation therapy (Cruz and Matushansky 2012, de Thé 2018).

A wide range of mathematical models have been powerful in highlighting the sometimes undercover role of cancer stem cells (see e.g. Michor 2008 and references therein). We here consider a minimal view of the accepted modeling of tissue architecture as a set of hierarchically organized cancer subpopulations (Michor et al. 2005, Dingli et al. 2007, Solé et al 2008, Figure 4). Following the previous approximations, we here add a seeding CSC compartment, *S*

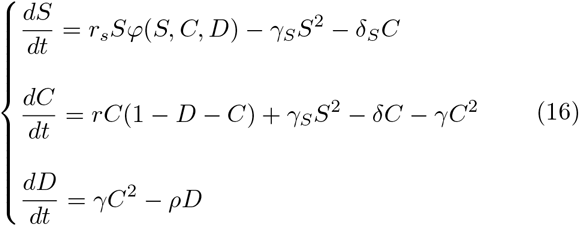

**FIG. 4:**
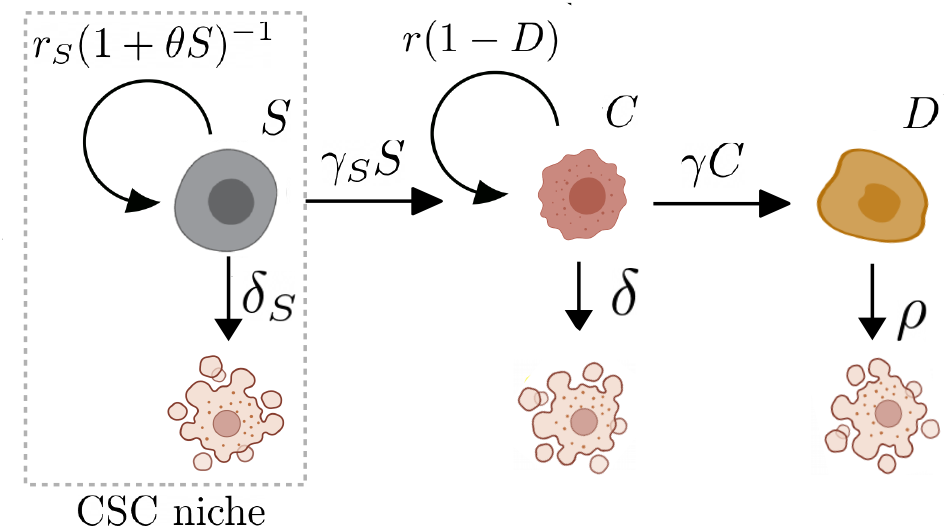
Tissue architecture and the ecology of tumor differentiation therapy. The minimal hierarchical model involves a cancer stem cell compartment *S* that replicates under the constraints of a well-separated niche, dies under cytotoxic therapy with diminished effectivity (*δ_S_* ⪡ *δ*) and seeds a progenitor cancer population *C*.

Here *C* indicates the progenitor compartment, that differentiates following Self-regulated mechanisms and feels the crowding (spatial) effects of these terminal cells *D*. This population, in turn, is seeded by a particular CSC compartment *S*, that replicates at rate *r_S_*. The effect of chemotherapy is captured by *δ_S_*, which is in generally much smaller than that of non-stem cancer cells due to plasticity or quiescence potential of CSC. Differentiation rate into the general progenitor compartment is captured by *γ_S_* and in a first approximation is believed to follow autocontrol regulation as in (Vainstein et al. 2012). Replication is again constrained by how populations occupy space, here *φ*(*S, C, D*). Stem cells (and CSCs in particular) are known to inhabit in a well-differentiated spatial niche (Plaks et al. 2015, Voog and Jones 2010), such as the bone marrow for hematopoietic stem cells (Addams and Scaden 2006). This indicates that, most often, CSC replication –in contrast with differentiation– might not be regulated by the density of the tumor populations. Here we have *φ*(*S, C, D*) = *φ*(*S*).

Furthermore, crowding effects observed in the bone marrow, together with niche and carrying capacity modulation (Dingli and Michor 2006, Gerlee and Anderson 2015), indicate that a logistic growth model with absolute saturation (*r*(*C* = *K*) = 0) might not be accurate to describe CSC plasticity. A more general saturating function *φ* considers an adimensional CSC sensitivity to crowding *θ* (see Dingli and Michor 2006). In our case, the model is written as

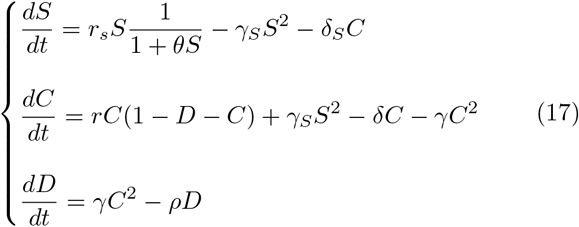

with the particularity that CSC replication does not totally stop, rather it slows down as the CSC niche becomes populated. This crowding effect cannot be correctly captured by cellular automata models studying DTH (such as Vainstein et al. 2012), as the number of neighbouring cells is considered constant. What is, once again, the effect of space in DTH? Could this CSC niche-specificity explain why some cancers are resistant to differentiating agents?

It can be seen how DTH, even if considered in a totally symmetric fashion where *γ_S_* = *γ*, does not have the same effect in a hierarchical tumor if CSCs inhabit a different niche (Fig. 3a). In particular, the model shows that, for low differentiation rates, the general cancer population needs a similar time until eradication as with the logistic growth model (figure 5a, red curves). However, as *γ* increases, not only CSCs residing in a different niche, but also the rest of the tumor becomes harder to eradicate (black curves). The model therefore captures how stem cell resistance to differentiation approaches not only resides in cellular plasticity (Foo et al 2009, Meacham and Morrison 2013), but also in the role of habitat ecology in niche construction and independence (Addams and Scaden 2006).

**FIG. 5:**
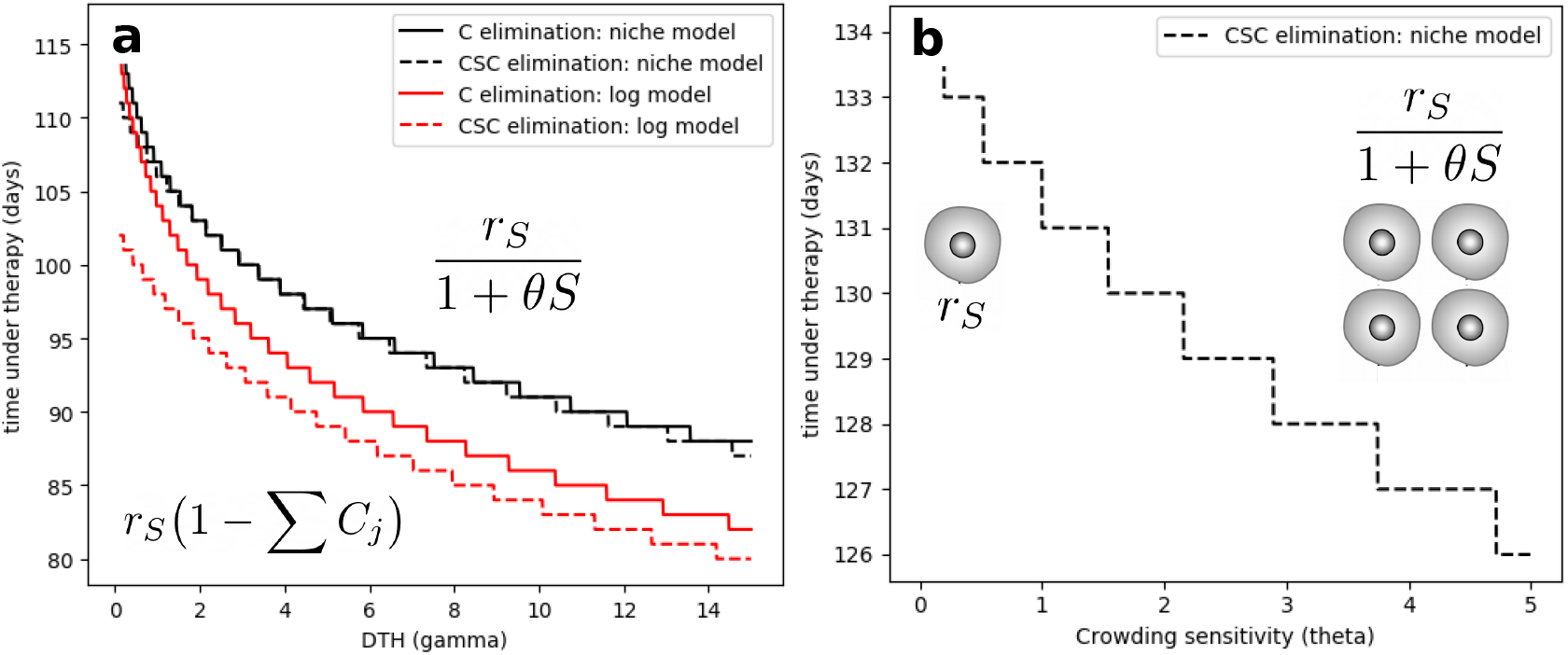
The role of DTH and spatial constraints in hierarchical tumor architectures. In **(a)**, the duration of therapy (for *δ_i_* > *r_i_*) until tumor eradication is computed as a function of DTH effectivity (*γ*). In particular, simulations compare the duration of therapy under a generic model (red) and a model that captures how CSCs inhabit a specific niche, and therefore do not feel the spatial effects of an increasingly differentiated habitat. In **(b)**, the effect of crowding sensitivity (*θ*) in the CSC model is studied. It is easily seen how an increasing sensitivity to surrounding cells results in reduced therapy duration.

Along with CSCs death and differentiation, sensitivity to CSC density *θ* also plays a role in tumor extinction. Increasing *θ* results in a linear decrease in the time of tumor eradication under chemotherapy (Figure 5b). This result, together with the overall indications of this work regarding the role of spatial effects of DTH, supports the application of *niche therapy* for CSCs (Plaks et al. 2015). Previous efforts had already considered the possibility of disrupting the CSC niche, such as through VEGF inhibition reducing blood vessel production (Calabrese et al. 2007). Together with this, our results propose that CSCs become weaker to DTH when increasing their sensitivity to the rest of tumor populations by physical disruption of their niche.

## IV. DISCUSSION

In this paper we have shown how ecological models of habitat loss can shed light into several aspects of differentiation therapy in cancer. This is done by using habitat loss as a surrogate of differentiated patches, while an independent extinction term is matched by the effects of cytotoxic therapy. Early models of ecological decline due to habitat loss show that a well-defined threshold exist: once a given critical loss is present, no viable populations are allowed, despite that some amount of habitat is still present (Levins 1969, Bascompte and Solé 1996). Within the cancer context, when a critical amount of cancer cells have been differentiated, remission results in a similar fashion, provided that standard cytotoxic therapy is also present as seen in the clinics of APL (de Th 2018). However, the dynamical nature of differentiation implies that not a sharp threshold, but rather a progression towards cancer eradication result from DTH. In order to test the generality of the approximation, both an homogeneous metapopulation model and an extension considering the specificity of a cancer stem cell compartment and its niche have been explored. The models consistently explain several qualitative observations concerning the impact of DTH.

On the one hand, approaching differentiation as an ecological process for a simple population and its terminally differentiated surrogate can predict interesting dynamics in genetically simple cancers such as APL where DTH has been successful. Our model predicts a well defined threshold for the amount of differentiated habitat, beyond which a malignant population is not able to progress. The fact that eradication with differentiating agents is totally dependent on cytotoxic therapy is consistent with studies on DTH for leukemia, where arsenic, that triggers p-53 driven senescence as well as differentiation, is effective as a single-agent therapy, while other agents might require combined cytotoxicity. Further learning from the single population model indicate that DTH becomes much more effective than chemotherapy for small, growing tumors away from the carrying capacity of their tissue. This result opens novel questions on the role of DTH as an early therapeutic scheme.

An extension of the ecological model introduces a minimal architecture to understand the possible role of a cancer stem cell niche in sensitivity to DTH. In particular, we aim at understanding how niche specificity in CSCs might be playing a role in resistance to DTH agents for certain tumor types. The model indicates how a hierarchical tumor seeded by a CSC population inhabiting a separated niche might be much more difficult to eradicate through DTH than in well-mixed approaches. This indicates that not only the cellular characteristics of cancer stem cells, but also the ecological interactions building their niche (namely, crowding and spatial competition) might be key in understanding their resistance to this kind of therapeutic approaches.

A large body of literature concerning mathematical modeling of cancer tissue hierarchies has grown in the course of recent years (Foley and Mackey 2008), from the understanding of cell fate decisions (Sun et al. 2015), the molecular basis of cytokine and division dynamics (Stiehl and Marciniak-Czochra 2012, Stiehl et al. 2015) or the study of control networks regulating CSC differentiation (Yang et al. 2017, Komarova et al. 2018), among many others. Despite using a similar fundamental background, our approach using the learning of habitat ecology is able to mathematically describe the phenomenological role of spatial interactions and how these, and the anatomical niches, might result in success or failure of DTH combined with other agents. Furthermore, the role of spatial distribution of CSC-niche interactions, and the possibility of disrupting the barriers of this distribution, appears as a possible improvement to the cytotoxic+DTH combination approach when cancer stem cells are in place.

Several shortcomings, potential extensions and implications of this work can be outlined. First of all, the model involves the most minimal set of rules and assumptions, sacrificing the details of the population description in favor of an ecological picture that can be intuitively interpreted. Real tumours include several layers of complexity, such as cell-cell interactions or micro-environmental cues that we have captured through phenomenological modeling alone. Moreover, models of habitat loss including noise reveal the importance of considering several sources of disturbance, from demographic stochasticity to large catastrophes (Casafrandi and Gatto 2002).

Further exploration would require considering, for example, the heterogeneous spatial organization and its Impact on tumor growth (Sottoriva et al 2010). However, the need for a more realistic model does not invalidate the key findings of our study. In fact, a similar criticism could be raised in relation to the simplicity of habitat loss models derived from Levins equation. Despite their minimalism, the simplest models (also ignoring the details of species-specific metabolic or physiologic features) have been extremely valuable in understanding the problem as well as how to prevent its consequences (Lande 1988, Hanski 2011).

## Acknowledgments

The authors thank the Complex Systems Lab members for fruitful discussions. Special thanks to Alvah Bassie for his inspiring ideas. This work was supported by the Boín Foundation by Banco Santander through its Santander Universities Global Division, the Spanish Ministry of Economy and Competitiveness, grant FIS2016-77447-R MINECO/AEI/FEDER, an AGAUR FI 2018 grant, and the Santa Fe Institute where most of this work was done.

